# Synthetic datasets and community tools for the rapid testing of ecological hypotheses

**DOI:** 10.1101/021402

**Authors:** Timothée Poisot, Dominique Gravel, Shawn Leroux, Spencer A. Wood, Marie-Josée Fortin, Benjamin Baiser, Alyssa R. Cirtwill, Miguel B. Araújo, Daniel B. Stouffer

## Abstract

The increased availability of both open ecological data, and software to interact with it, allows the fast collection and integration of information at all spatial and taxonomic scales. This offers the opportunity to address macroecological questions in a cost-effective way. In this contribution, we illustrate this approach by forecasting the structure of a stream food web at the global scale. In so doing, we highlight the most salient issues needing to be addressed before this approach can be used with a high degree of confidence.

---

Ecologists are often asked to provide information and guidance to solve a variety of issues, across different scales. As part of the global biodiversity crisis, notable examples include predicting the consequences of the loss of trophic structure (Estes et al. 2011), rapid shifts in species distributions (Gilman et al. 2010), and increased anthropogenic stress on species and their environment. Most of these pressing issues require the integration of a variety of ecological data and information, spanning different geographical and environmental scales, to be properly addressed (Thuiller et al. 2013). Because of these requirements, relying solely on *de novo* sampling of the ecological systems of interests is not a viable solution on its own. Chiefly, there are no global funding mechanisms available to finance systematic sampling of biological data, and the spatial and temporal scales required to acquire meaningful data on biodiversity change are such that it would take a long time before realistic data would be available to support the decision process. While that data collection must continue, we propose that there are a large number of macroecological questions that could be addressed without additional data or with data acquired at minimal cost, by making use of open data and community-developed software and platforms.

Existing datasets can, to an increasing extent, be used to *build* new datasets (henceforth *synthetic* datasets, since they represent the synthesis of several types of data). There are several parallel advances that make this approach possible. First, the volume of data on ecological systems that are available *openly* increases on a daily basis. This includes point-occurrence data (as in GBIF or BISON), but also taxonomic knowledge (through ITIS, NCBI or EOL) or trait and interactions data. In fact, there is a vast (and arguably under-exploited) amount of ecological information, that is now available without having to contact and secure authorization from every contributor individually. Second, these data are often available in a *programmatic* way; as opposed to manually visiting data repositories, and downloading or copy-and-pasting datasets, several software packages offer the opportunity to query these databases automatically, considerably speeding up the data collection process. As opposed to manual collection, identification, and maintenance of datasets, most of these services implement web APIs (Application Programming Interface, *i.e.* services that allow users to query and/or upload data in a standard format). These services can be queried, either once or on a regular basis, to retrieve records with the desired properties. This ensures that the process is repeatable, testable, transparent, and (as long as the code is properly written) nearly error proof. Finally, most of the heavy lifting for these tasks can be done through a *burgeoning ecosystem of packages and software* that handles query formatting, data retrieval, and associated tasks, all the while exposing simple interfaces to researchers. None of these are *new* data, in the sense that these collections represent the aggregation of thousands of ecological studies; the originality lies in the ability to query, aggregate, curate, and use these data consistently and in a new way using open solutions.

Hypothesis testing for large-scale systems is inherently limited by the availability of suitable datasets – most data collection results in small scale, local data, and it is not always clear how these can be used at more global scales. Perhaps as a result, developments in macroecology have primarily been driven by a search for patterns that are very broad both in scale and nature (Keith et al. 2012, Becknell et al. 2015). While it is obvious that collecting exhaustive data at scales that are large enough to be relevant can be an insurmountable effort (because of the monetary, time, and human costs needed), we suggest that macroecologists could, in parallel, build on existing databases, and aggregate them in a way that allows direct testing of proposals stemming from theory. To us, this opens no less than a new way for ecologists to ask critical research questions, spanning from the local to the global, and from the organismal to the ecosystemic, scales. Here, we (i) outline approaches for integrating data from a variety of sources (both in terms of provenance, and type of ecological information), (ii) identify technical bottlenecks, (iii) discuss issues related to scientific ethics and best practice, and (iv) provide clear recommendations moving forward with these approaches at larger scales. Although we illustrate the principles and proposed approaches with a real-life example, the objective of this paper is to highlight the way different tools can be integrated in a single study, and to discuss the current limitations of this approach. This approach can, for example, prove particularly fruitful if it allows researchers to either offer new interpretation of well-described macroecological relationships, or to provide tests of hypotheses suggested by theoretical studies (Levin 2012).

## 1 An illustrative case-study

Food-web data, that is the determination of trophic interactions among species, are notoriously difficult to collect. The usual approach is to assemble literature data, expert knowledge, and additional information coming from field work, either as direct observation of feeding events or through gut-content analysis. Because of these technical constraints, food-web data are most often assembled based on sampling in a single location. Assessments of food web structure over space may therefore require comparisons of communities composed of different taxa. As a consequence, most food web properties over large (continental, global) spatial extents remain undocumented. For example, what is the relationship between latitude and connectance (the density of feeding interactions)? One possible way to approach this question is to collect data from different localities, and document the relationship between latitude and connectance through regressions. The approach we illustrate uses broad-scale data integration to forecast the structure of a single system at the global scale (Pellissier et al. 2013). We are interested in predicting the structure of a pine-marsh food web, worldwide.

### 1.1 Interactions data

The food-web data were taken from Thompson et al. (2012), as made available in the IWDB database (https://www.nceas.ucsb.edu/interactionweb/html/thomps_towns.html) – starting from the Martins dataset (stream food web from a pine forest in Maine). Wetlands and other freshwater ecosystems are critically endangered and serve as a home to a host of endemic biodiversity (Fensham et al. 2011, Minckley et al. 2013). Stream food webs in particular are important because they couple terrestrial and aquatic communities and ensure the maintenance of ecosystem services such as freshwater quality and flood regulation. Anthropogenic pressure on wetlands makes them particularly threatened. They represent a prime example of ecosystems for which data-driven prediction can be used to generate scenarios at a temporal scale relevant for conservation decisions, and at a faster rate than sampling allows.

The data from the original food web had 105 nodes, including vague denominations like *Unidentified detritus* or *Terrestrial invertebrates.* First, we aggregated all nodes to the *genus* level. Due to the high level of structure in trophic interactions emerging from taxonomic rank alone (Eklof et al. 2011, Stouffer et al. 2012, Eklöf and Stouffer 2015), aggregating to the genus level has the double advantage of (i) removing ambiguities on the identification of species and (ii) allowing integration of data when any two species from given genera interact. Second, we removed all nodes that were not identified (`Unidentified` or `Unknown` in the original data). The cleaned network documented 227 interactions, between 80 genera. We then used the name-checking functions from the `taxize` package (Chamberlain and Szöcs 2013) to perform the following steps. First, all names were resolved, and one of the following was applied: valid names were conserved, invalid names with a close replacement were corrected, and invalid names with no replacement were removed. In most situations, invalid names were typos in the spelling of valid ones. After this step, 74 genera with 189 interactions remained, representing a high quality genus-level food-web from the original sampling.

Because this food web was sampled *locally,* there is the possibility that interactions between genera are not reported; either because species from these genera do not interact or do not co-occur in the sampling location, or because of spatial mismatches between genus occurrence and sampling. To circumvent this, we queried the *GLOBI* database (Poelen et al. 2014) for each genus name, and retrieved all *feeding* interactions; this includes taxa from the original dataset, but also taxa that establish interactions with them even though these were not observed in the original sample. For all *new* genera retrieved through this method, we also retrieved their interactions with genera already in the network. The inflated network (original data plus data from *GLOBI)* has 368 genera, and a total of 4796 interactions between them.

As a final step, we queried the GBIF taxonomic rank database with each of these (tentatively) genera names. Every tentative genus that was either not found, or whose taxonomic level was not *genus*, was removed from the network.

The code to reproduce this analysis is in the `1_get_data.r` suppl. file.

It should be noted that this analysis relies on *databases,* and a vast majority of information is confined to the primary literature. While it is possible to do manual literature surveys (e.g. Strong and Leroux 2014), this task becomes daunting for large number of species. Initiatives like text-mining (Milani et al. 2012) will speed up the rate at which we can recover interactions data from the literature – if publishers allow researchers to mine the literature they create.

### 1.2 Occurrence data and filtering

For each genus, we retrieved the known occurrences (approx. 2 × 10^5^) from GBIF and BISON. Because the ultimate goal is to perform spatial modeling of the structure of the network, we removed genera for which fewer than 100 occurrences in the entire dataset. This stringent filter enables us (i) to maintain high predictive powers for SDMs, and (ii) to work on the genera for which we have “high-quality” data. The cleaned food web had a total of 134 genera and 782 interactions, for 118269 presences. Given the curated publicly available data, it represents the current best description of feeding interactions between species of this ecosystem. A visual depiction of the network is given in Figure 1.

**Figure 1:**
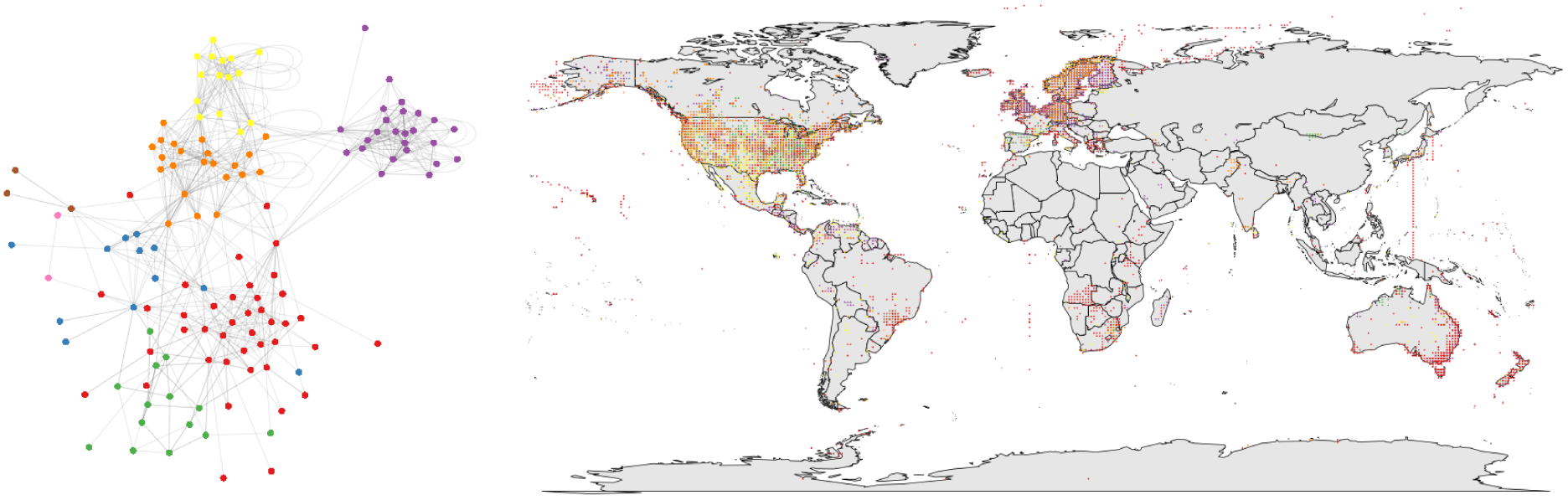
Visual representation of the initial data. On the left, we show the food web (original data and interactions from GLOBI), with genera forming modules (clusters of densely connected nodes) in different colors. On the right, we show the occurrence data where each dot represents one observation from BISON and GBIF (again color coded by module)

On its own, the fact that filtering for genera with over 100 records reduced the sample size from 368 genera to 134 indicates the importance of the deposition of all observations in public databases. This is because the analysis we present here, although cost-effective and enabling rapid evaluation of different scenarios, is only as good as the underlying data. Since most modeling tools require a minimal sample size in order to achieve acceptable accuracy, concerted efforts by the community and funding agencies to ensure that the minimal amount of data is deposited upon publication or acquisition is needed. It must also be noted that the threshold of 100 occurrences is an arbitrary one.

The approach is amenable to sensitivity analysis, and indeed this will be a crucial component of future analyses. A taxon can have less observations than the threshold either because of under-sampling or under-reporting, or because it is naturally rare. In the context of food webs, species higher-up the food chain can be less common than primary producers. To which extent these relationships between, *e.g.,* trophic position and rarity, can influence the predictions, will have to receive attention.

The code to reproduce this analysis is in the `1_get_data.r` suppl. file.

### 1.3 Species Distribution Model

For each species in this subset of data, we retrieved the nineteen `bioclim` variables (Hijmans et al. 2005), with a resolution of 5 arc-minutes. This enabled us to build climatic envelope models, using *biolcim,* for each species. These models tend to be more conservative than alternate modeling strategies, in that they predict smaller range sizes (Hijmans and Graham 2006), but they also perform well overall for presence-only data (Elith et al. 2006, Elith and Graham 2009). The output of these models is, for species *i*, the probability of an observation P(*i*) within each pixel. We appreciate that this is a coarse analysis, but its purpose is to highlight how to combine different data. A discussion of the limitations of this approach is given below.

The code to reproduce this analysis is in the `2_get_sdm.r` suppl. file.

### 1.4 Assembly

For every interactions in the food web, we estimated the probability of it being observed in each pixel as the product of the probabilities of observing each species on its own: P(*L_ij_*) ∝ P(*i*)P(*j*). This resulted in one LDM (“link distribution model”) for each interaction. It should be noted that co-occurrence is considered to be entirely neutral, in that we assume that the probability that two species co-occur is independent (*i.e.* a predator is not more likely to be present if there are, or are not, potential prey). We also assume no variability in interactions, as in Havens (2015). It is likely that, in addition to their occurrence, species co-occurrences and interactions (Poisot et al. 2015) are affected by climate. Whether or not these constitute acceptable assumptions has to be decided for each study.

The code to reproduce this analysis is in the `3_get_ldm.r` suppl. file.

Based on this information, we generated example illustrations (using `4_draw_figures.r` – Figure 2). The system is characterized, at the world-wide scale, by an increased number of genera *and* interactions in temperate areas, with diversity and interaction hotspots in Western Europe, North-East and South-Atlantic America, and the western coasts of New Zealand and Australia – this is clearly symmetrical along the equator. Network structure, here measured by network connectance, follows a different trend than genera richness or interactions do. Connectance is stable along the gradient, but declines at extreme latitudes (Figure 2B).

**Figure 2:**
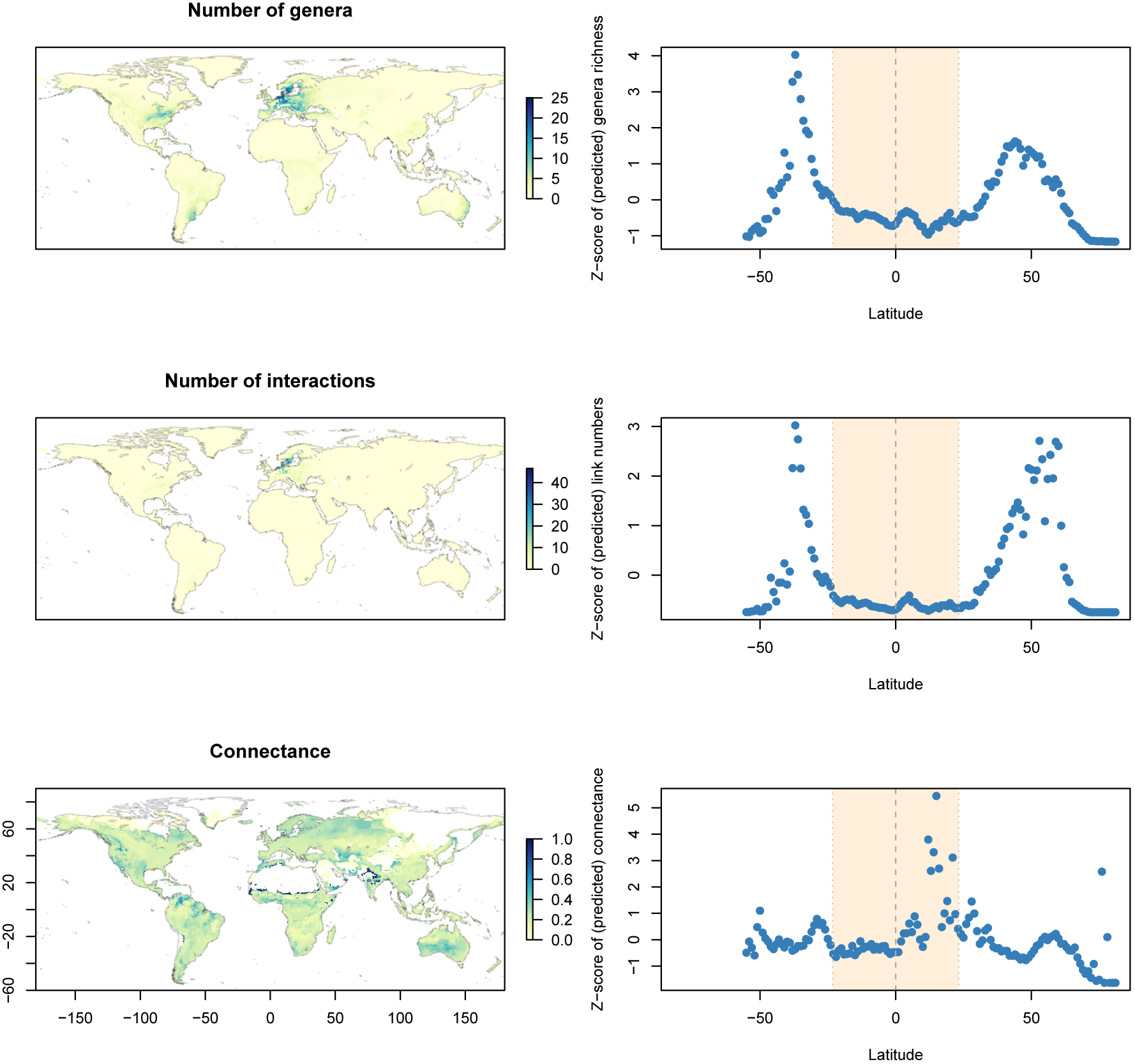
Maps for the number of genera, number of interactions, and connectance in the assembled networks (on the left) as well as their underlying relationship with latitude (on the right). The tropics are shaded in light yellow. The average value of each output has been (i) averaged across latitudes and (ii) z-score transformed; this emphasizes variations across the gradient as opposed to absolute values (which is a more conservative way of looking at the results since the predictions are qualitative).

## 2 Challenges moving forward

The example provided illustrates the promises of data-driven approaches. It builds on new data availability, new statistical and computational tools, and new ways to integrate both. Most importantly, it allows us to use “classical” ecological data in a resolutely novel way, thus presenting an important opportunity to bridge a gap between field-based and theory-based macroecological research. But as with every methodological advancement, comes a number of challenges and limitations. Here we discuss a few we believe are important. In doing so, we hope to define these issues and emphasize that each of them, on their own, should be the subject of further discourse.

### 2.1 Attribution stacking and intellectual provenance

The merging of large databases has already created a conflict of how to properly attribute data provenance (Carroll 2015). Here there are at least two core issues that will require community consultation in order to be resolved. First, *what is the proper mode of attribution when a very large volume of data are aggregated*? Second, *what should be the intellectual property of the synthetic dataset*? Currently, citations (whether to articles or datasets) are only counted when they are part of the main text. The simple example outlined here relies on well over a thousand references, and it makes little sense to expect that they would be provided in the main text (nor do we expect any journal to accept a manuscript with over a hundred references or so, with rare exceptions). One intermediate solution would be to collate these references in a supplement, but it is unclear that these would be counted (Seeber 2008), and therefore contribute to the *impact* of each individual dataset (and hence, collector; Kueffer et al. 2011). This is a problem that we argue is best solved by publishers; proper attribution and credit is key to provide incentives to data release (Kenall et al. 2014, Whelan et al. 2014, Pronk et al. 2015). As citations are currently the “currency” of scientific impact, publishers have a responsibility not only to ensure that data are available (which many already do), but that they are recognized; data citation, no matter how many data are cited, is a way to achieve this goal. The synthetic dataset, on the other hand, can reasonably be understood as a novel product; there is technical and intellectual effort involved in producing it, and although it is a derivative work, we would encourage authors to deposit it anew. Nevertheless, we would like to see a more meaningful dialogue between editors, publishers, and authors, to determine how the citation of thousands of datasets ought to be handled across the editorial process.

### 2.2 Sharing of code and analysis pipeline

Ideally, authors should release their analysis *pipeline* (that is, the series of steps, represented by code, needed to reproduce the analysis starting from a new dataset) in addition to the data and explanation of the steps. The pipeline can take the form of a `makefile` (which allows one to generate the results, from the raw data, without human intervention), or be all of the relevant code that allows to re-generate the figures and results. For example, we have released all of the R code that was used to generate the figures at https://zenodo.org/record/31975. Sharing the analysis pipeline has several advantages. First, it is a first step towards ensuring the quality of analyses, since reviewers can (and should reasonably be expected to) look at the source code. Second, it provides a *template* for future analyses – instead of re-developing the pipeline from scratch, authors can re-use (and acknowledge) the previous code base and build on it. Finally, it helps identify areas of future improvement. The development of software should primarily aim to make the work of researchers easier. Looking at commonalities in the analytical pipelines for which no ready-made solutions exists will be a great way to influence priorities in software development. Properly citing and reviewing computer code is still an issue, because software evolves whereas papers remain (for now) frozen in the state where they were published. Being more careful with citation, notably by including version number (White 2015) or using unique identifiers (Poisot 2015), will help longterm reproducibility.

### 2.3 Computational literacy

This approach hardly qualifies as *big data*; nevertheless, it relies on the management and integration of a large volume of heterogeneous information, both qualitatively larger than the current “norm”. The first challenge is being able to *manage* these data; it requires data management skills that are not usually needed when the scale of the dataset is small, and, fallible though the process may be, when data can reasonably be inspected manually. The second challenge is being able to *manipulate* these data; even within the context of this simple use-case, the data do not fit in the memory of R (arguably the most commonly known and used software in ecology) without some adjustments. Once these issues were overcome, running the analysis involved a few hours worth of computation time. Computational approaches are going to become increasingly common in ecology (Hampton et al. 2012, 2013), and are identified by the community as both in-demand skills and as not receiving enough attention in current ecological curricula (Barraquand et al. 2014). It seems that efforts should be allocated to raise the computational literacy of ecologists, and recognize that there is value in the diversity of tools one can use to carry out more demanding studies. For example, both `Python` and `Julia` are equally as user friendly as `R` while also being more powerful and better suited for computationally- or memory-intensive analyses.

### 2.4 Standards and best practices

In conducting this analysis, we noticed that a common issue was the identification of species and genera. All of these datasets were deposited by individual scientists; whether we like it or not, individuals are prone to failure in a very different way than the “Garbage in, garbage out” idea that applies to computer programs. Using tools such as `taxize` (Chamberlain and Szöcs 2013) can allow us to resolve a few of the uncertainties, yet this must be done every time the data are queried and requires the end user to make educated guesses as to what the “true” identity of the species is. These limitations can be overcome, on two conditions. Database maintainers should implement automated curation of the data for which they are stewards, and identify potential mistakes and correct them upstream, so that users download high-quality, high-reliability data. Data contributors should rely more extensively on biodiversity identifiers (such as TSN, GBIF, NCBI Taxonomy ID, etc.), to make sure that even when there are typos in the species name, they can be matched across datasets. Constructing this dataset highlighted a fundamental issue: the rate-limiting step is rarely the availability of appropriate tools or platforms, but instead it is the adoption of common standards and the publication of data in a way that conforms to them. In addition, Maldonado et al. (2015) emphasize that point-occurence data are not always properly reported – for example, the center of a country or region can be used when no other information is known; this requires an improved dialogue between data collectors and data curators, to highlight which practices have the highest risk of biasing future analyses.

### 2.5 Propagation of error

There are always caveats to using synthetic datasets. First, the extent to which each component dataset is adequately sampled is unknown (although there exist ways to assess the overall representativeness of the assembled dataset; Schmill et al. (2014)). This can create gaps in the information that is available when all component datasets are being merged. Second, because it is unlikely that all component datasets were acquired using reconcilable standards and protocol, it is likely that much of the quantitative information needs be discarded, and therefore the conservative position is to do qualitative analyses only. Although these have to be kept in mind, we do not think they are so sufficient as to prevent use and evaluation of the approach we suggest. For one thing, as we illustrate, at large spatial and organizational scales, coarse-grained analyses are still able to pick up qualitative differences in community structure. Second, most emergent properties are relatively insensitive to fine-scale error; for example, Gravel et al. (2013) show that even though a simple statistical model of food-web structure mispredicts some individual interactions, it produces communities with realistic emergent properties. Which level of error is acceptable needs to be determined for each application, but we argue that the use of synthetic datasets is a particularly cost- and time-effective approach for broad-scale description of community-level measures.

## 3 Conclusion – why not?

In light of the current limitations and challenges, one might be tempted to question the ultimate validity and utility of this approach. Yet there are several strong arguments, that should not be overlooked, in favor of its use. As we demonstrate with this example, synthetic datasets allow us to rapidly generate qualitative predictions at large scales. These can, for example, serve as a basis to forecast the effect of scenarios of climate change on community properties (Albouy et al. 2012, 2014). Perhaps more importantly, synthetic datasets will be extremely efficient at identifying gaps in our knowledge of biological systems: either because there is high uncertainty or sensitivity to choices in the model output, or because there is no available information to incorporate in these models. By building these datasets, it will be easier to assess the extent of our knowledge of biodiversity, and to identify areas or taxa of higher priority for sampling. For this reason, using synthetic datasets is *not* a call to do less field-based science. Quite the contrary: in addition to highlighting areas of high uncertainty, synthetic datasets provide *predictions* that require field-based validation. Only through this feedback can we build enough confidence in this approach to apply it for more ambitious questions, or disqualify it altogether. Meanwhile, the use of synthetic datasets will necessitate the development of both statistical methodology and software; this is one of the required steps towards real-time use and analysis of ecological data (Antonelli et al. 2014). We appreciate that this approach currently comes with some limitations – they are unlikely to be overcome except with increased use, testing, and validation. Since the community already built effective and user-friendly databases and tools, there is very little cost (both in time and in funding) in trying these methods and there is also the promise of great potential.

## Acknowledgments

This work was funded in part through a grant from the Canadian Institute of Ecology and Evolution. TP was funded by a Starting grant from the Université de Montréal, and a NSERC Discovery Grant. SL was funded by a NSERC Discovery Grant. DBS acknowledges a Marsden Fund Fast-Start grant (UOC-1101) and Rutherford Discovery Fellowship, both administered by the Royal Society of New Zealand. We thank Kévin Cazelles for constructive comments on the manuscript. We thank Anne Bruneau and Andrew Gonzalez for organizing the workshop at which this approach was first discussed. We thank Ross Mounce and one anonymous reviewer for comments on the manuscript.

## References

Albouy, C. et al. 2012. Projected climate change and the changing biogeography of coastal Mediterranean fishes (P Pearman, Ed.). - J. Biogeogr. 40: 545–547.

Albouy, C. et al. 2014. From projected species distribution to food-web structure under climate change. - Glob Change Biol 20: 730–741.

Antonelli, A. et al. 2014. SUPERSMART: ecology and evolution in the era of big data. 2

Barraquand, F. et al. 2014. Lack of quantitative training among early-career ecologists: a survey of the problem and potential solutions. - PeerJ 2: e285.

Becknell, J. M. et al. 2015. Assessing Interactions Among Changing Climate, Management, and Disturbance in Forests: A Macrosystems Approach. - BioScience 65: 263–274.

Carroll, M. W. 2015. Sharing Research Data and Intellectual Property Law: A Primer. - PLoS Biol 13: e1002235.

Chamberlain, S. A. and Szöcs, E. 2013. taxize: taxonomic search and retrieval in R. - F1000Research in press.

Eklof, A. et al. 2011. Relevance of evolutionary history for food web structure. - Proc. R. Soc. B Biol. Sci. 279: 1588–1596.

Eklöf, A. and Stouffer, D. B. 2015. The phylogenetic component of food web structure and intervality. - Theor Ecol in press.

Elith, J. and Graham, C. H. 2009. Do they? How do they? WHY do they differ? On finding reasons for differing performances of species distribution models. - Ecography 32: 66–77.

Elith, J. et al. 2006. Novel methods improve prediction of species’ distributions from occurrence data. - Ecography 29: 129–151.

Estes, J. A. et al. 2011. Trophic Downgrading of Planet Earth. - Science 333: 301–306.

Fensham, R. J. et al. 2011. Four desert waters: setting arid zone wetland conservation priorities through understanding patterns of endemism. - Biol. Conserv. 144: 2459–2467.

Gilman, S. E. et al. 2010. A framework for community interactions under climate change. - Trends Ecol. Evol. 25: 325–331.

Gravel, D. et al. 2013. Inferring food web structure from predatorprey body size relationships. - Methods Ecol Evol 4: 1083–1090.

Hampton, S. E. et al. 2012. Ecological data in the Information Age. - Frontiers in Ecology and the Environment 10: 59–59.

Hampton, S. E. et al. 2013. Big data and the future of ecology. - Frontiers in Ecology and the Environment 11: 156–162.

Havens, K. 2015. Scale and structure in natural food webs. - Science 257: 1107–1109.

Hijmans, R. J. and Graham, C. H. 2006. The ability of climate envelope models to predict the effect of climate change on species distributions. - Glob. Change Biol. 12: 2272–2281.

Hijmans, R. J. et al. 2005. Very high resolution interpolated climate surfaces for global land areas. – Int. J. Climatol. 25: 1965–1978.

Keith, S. A. et al. 2012. What is macroecology? - Biology Letters: rsbl20120672.

Kenall, A. et al. 2014. An open future for ecological and evolutionary data? - BMC Ecology 14: 10.

Kueffer, C. et al. 2011. Fame, glory and neglect in meta-analyses. - Trends in Ecology & Evolution 26: 493–494.

Levin, S. A. 2012. Towards the marriage of theory and data. - Interface Focus 2: 141–143.

Maldonado, C. et al. 2015. Estimating species diversity and distribution in the era of Big Data: to what extent can we trust public databases? - Global Ecology and Biogeography: n/a-n/a.

Milani, G. A. et al. 2012. Machine Learning and Text Mining of Trophic Links. - 2012 11th International Conference on Machine Learning and Applications

Minckley, T. A. et al. 2013. The relevance of wetland conservation in arid regions: a re-examination of vanishing communities in the American Southwest. - J. Arid Environ. 88: 213–221.

Pellissier, L. et al. 2013. Combining food web and species distribution models for improved community projections. - Ecol. Evol. 3: 4572–4583.

Poelen, J. H. et al. 2014. Global Biotic Interactions: An open infrastructure to share and analyze species-interaction datasets. - Ecological Informatics 24: 148–159.

Poisot, T. 2015. Best publishing practices to improve user confidence in scientific software. - Ideas in Ecology & Evolution in press.

Poisot, T. et al. 2015. Beyond species: why ecological interaction networks vary through space and time. - Oikos 124: 243–251.

Pronk, T. E. et al. 2015. A game theoretic analysis of research data sharing. - PeerJ 3: e1242.

Schmill, M. D. et al. 2014. GLOBE: Analytics for Assessing Global Representativeness. - 2014 Fifth International Conference on Computing for Geospatial Research and Application in press.

Seeber, F. 2008. Citations in supplementary information are invisible. - Nature 451: 887–887.

Stouffer, D. B. et al. 2012. Evolutionary Conservation of Species’ Roles in Food Webs. - Science 335: 1489–1492.

Strong, J. S. and Leroux, S. J. 2014. Impact of Non-Native Terrestrial Mammals on the Structure of the Terrestrial Mammal Food Web of Newfoundland, Canada. - PloS One 9: e106264.

Thompson, R. M. et al. 2012. Food webs: reconciling the structure and function of biodiversity. - Trends Ecol. Evol.: 1–9.

Thuiller, W. et al. 2013. A road map for integrating eco-evolutionary processes into biodiversity models. - Ecol. Lett. 16: 94–105.

Whelan, A. M. et al. 2014. Editorial. - Ecological Applications 24: 1–2.

White, E. P. 2015. Some thoughts on best publishing practices for scientific software. - Ideas in Ecology & Evolution in press.

